# Directed Evolution of a *Pantoea* Strain for Antibiotic Production via Interspecies Competition: A Reproducible Experimental Approach

**DOI:** 10.1101/2025.06.13.659473

**Authors:** Jaroslav Pavelka, Kristýna Jánošková, Karolína Hlinková, Eliška Horová, Jaroslav Pavelka, Radek Šíma

**Affiliations:** Centre for Biology, Geosciences and the Environment, Faculty of Education, University of West Bohemia, Univerzitní 22, 301 00, Pilsen, Czech Republic; 2nd Primary School Plzeň, Schwarzova 20, contributory organization, Schwarzova 20, 301 00 Plzeň, Czech Republic; Bioptic Laboratory, Mikulášské náměstí 4, 326 00, Plzeň, Czech Republic; Department of Pathology, Faculty of Medicine in Plzen, Charles University, and University Hospital, E. Beneše 13, 305 99 Plzeň, Czech Republic

**Keywords:** Antibiotics, bacterial competition, *Staphylococcus aureus*, *Pantoea*, *Acinetobacter baumannii*, *Streptococcus agalactiae*, *Escherichia coli*, *Anaplasma phagocytophilum*, *Pseudomonas aeruginosa*

## Abstract

A novel methodology for evolving antibiotic-producing bacterial strains is presented, based on interspecies competition and environmental pressure. Using a previously undescribed strain of the genus *Pantoea*, we established a breeding protocol in which gradual exposure to different bacterial pathogens at increasing temperatures resulted in the selective emergence of antibiotic-producing variants. The target pathogens included *Staphylococcus aureus*, *Acinetobacter baumannii*, *Streptococcus agalactiae*, *Escherichia coli*, and *Pseudomonas aeruginosa*. The competitive passages were performed over a range of temperatures, gradually selecting for strains capable of inhibiting these species. Antibiotic activity was confirmed via disk diffusion assays. Although the chemical identity of the active compounds remains to be determined, their biological effects are reproducible and strain-specific. The approach provides a low-cost, scalable method for the generation of new antibiotic-producing bacterial strains.

## Introduction

Antibiotics save lives around the world, yet the growing resistance of bacteria poses a major global health threat (1). To address this, new strategies for discovering antibacterial compounds are urgently needed. In 2024, the WHO published a list of antibiotic-resistant bacteria, underscoring the urgent need for the discovery of new antibiotics and alternative treatment strategies (2). While genomic mining and synthetic biology have proven powerful, they remain inaccessible to many laboratories due to cost or technical constraints. As an alternative, we present here a biological and scalable method to induce antibiotic production through evolutionary pressure in vitro.

It is well established that bacterial colonies can engage in competitive interactions, producing antibiotics to inhibit each other in the struggle for nutrients and ecological dominance (3). Notably, the production of such antimicrobial compounds is often triggered under stressful environmental conditions, which impose selective pressure and drive bacterial adaptation (3–4). A novel methodology for evolving antibiotic-producing bacterial strains is presented, based on interspecies competition and environmental pressure. Using a previously undescribed strain of the genus *Pantoea*, we established a breeding protocol in which gradual exposure to different bacterial pathogens at increasing temperatures resulted in the selective emergence of antibiotic-producing variants. The target pathogens included *Staphylococcus aureus*, *Acinetobacter baumannii*, *Streptococcus agalactiae*, *Escherichia coli*, and *Pseudomonas aeruginosa*. The competitive passages were performed over a range of temperatures, gradually selecting for strains capable of inhibiting these species. Antibiotic activity was confirmed via disk diffusion assays. While the precise chemical nature of the active substances has yet to be elucidated, their reproducible inhibitory effects and specificity to the producing strain have been consistently observed. This strategy offers an affordable and easily scalable pathway for evolving bacterial strains capable of synthesizing novel antibacterial agents.

Selective breeding in bacteria occurs rapidly, not only due to their short generation times— often measured in hours—but also because each generation produces millions of cells, increasing the likelihood of beneficial mutations. Moreover, bacteria possess the ability to exchange genetic material through horizontal gene transfer, particularly via plasmids. This mechanism is well known for its role in spreading antibiotic resistance among bacterial populations (5). However, here we assume that bacteria transfer genes for the production of new types of antibiotics.

The well-described and related bacterium *Pantoea agglomerans* (*Erwenia herbicola, Enterobacter agglomerans*) is a non-pathogenic microorganism that can produce the antibiotic pantocin. *P. agglomerans* inhibits the closely related bacterium *E. amylovora* by producing antibiotics against it. Pantocin is structurally a small peptide molecule. Two structures of this antibiotic have been found, pantocin A and pantocin B. Both antibiotics produced are capable of inhibiting histidine or arginine biosynthesis via transaminase (6–7) reported that *P. agglomerans* and its related *P. disperza* can, using cosmids pCPP702 and pCPP704, confer antibiotic-producing properties to other bacteria, such as *Escherichia coli*. The effect of pantocin is through the use of the tripeptide Ala-Gly-Gly, which is used by tripeptide transporters for transport into the cell (8).

Our primary aim was not to isolate a specific antibiotic molecule, but to validate a methodology for inducing stable inhibitory phenotypes that can be detected, replicated, and ultimately characterized. Our results demonstrate that this approach reliably generates strains with antibacterial activity, verified via disk diffusion assays, and adapted for activity at temperatures compatible with mammalian physiology.

## Material and methods

### Bacterial cultivation and selective breeding

Bacteria Z1*, S. aureus*, *S. agalactiae*, *A. baumannii*, *E. coli* and *P. aeruginosa* were used for bacterial cultivation. Bacteria Z1 classified in the genus *Pantoea* was isolated from bat droppings, probably from *Myotis myotis* at the Točník Castle site (Czech Republic) (Horová 2022) (6) (Fig 1). Other bacteria were obtained from The Czech Collection of Microorganisms (CCM), Masaryk University, Faculty of Science, Kamenice 5, building E25, Brno, Czech republic and the selected strains are listed under the following numbers: *S. aureus* CCM 4516, *S. agalactiae* CCM 6187, two strains *of A. baumanii* CCM no. 2355 and CCM no. 7031. We selected two types of bacteria from the WHO list against which current antibiotics have little effect. *S. agalactiae*, which is not on the WHO list, was also selected. These bacteria were chosen for their relative safety for healthy people. *Sacharomyces cerevisiae* was obtained from a common food distribution network in the Czech Republic, where it is sold in live form.

**Fig. 1.**
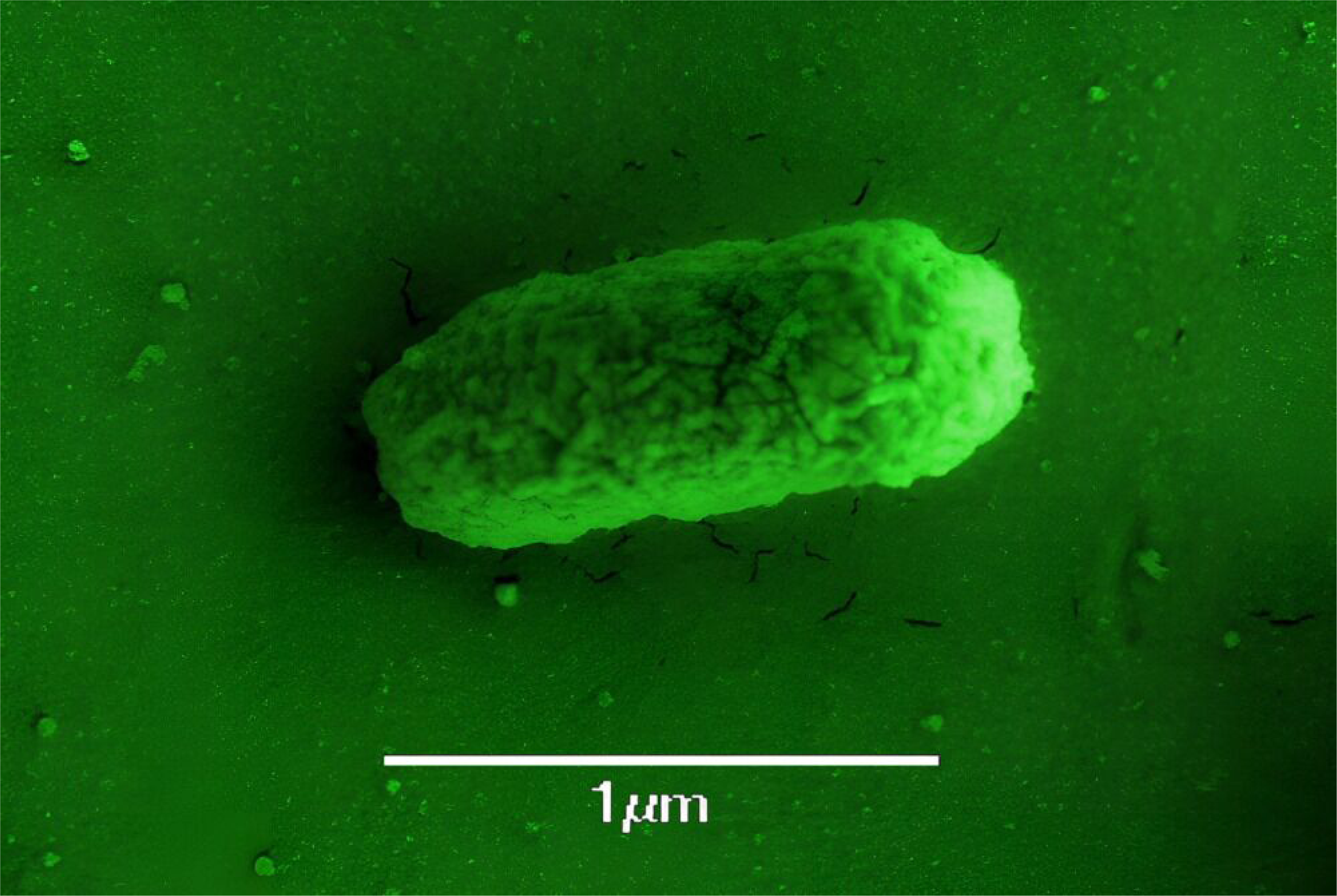
Electron microscope photograph of the bacterium Z1

Cultivation was carried out on LB (Luria-Bertani) agar plates of Petri dishes and in liquid medium. For the cultivation of *A. baumannii* strains and *P. aeruginosa*, LB medium was later mixed 1:1 with tryptone-soy agar. First, the bacteria were cultivated in liquid medium on a shaker for 24-48 hours. In the case of *E. coli* and *P. aeruginosa*, the competition took place only in Petri dishes. *S. aureus* against the antibiotic-producing bacterium Z1 was added. The same procedure was followed for Z1, *S. agalactiae A. baumannii, E. coli* and *P. aeruginosa*. Subsequently, the bacteria were plated together on agar plates of Petri dishes, spread with a stick and placed in a thermostat for 24-48 hours. When the bacteria grew on the Petri dishes, preparations (from the Petri dishes and from the liquid medium) were made for observation under an Olympus BX53 microscope with immersion oil at a magnification of 1000x. Throughout the contact of the two bacterial colonies, competitive interactions between bacteria occurred in the liquid medium and on the agar plates of Petri dishes. The goal was to obtain an antibiotic against *S. aureus*, *S. agalactiae, A. baumannii* and *P. aeruginosa* by breeding, and a partial goal was to breed the antibiotic to function at higher temperatures (with a limit of 38°C). Cultivation worked best on Petri dishes. Contamination in the form of unwanted bacteria was more likely to enter and remain in the liquid medium.

The cultivation of antibiotics using Z1 bacteria began at 12°C in a ratio of 1:1 (Table 1). At this temperature, Z1 bacteria produced the antibiotic best against *S. aureus*. We considered whether the bacteria was cold-loving and therefore chose a temperature of 12°C. Bacterial passage was used for the cultivation of antibiotics. Each passage was made up of different ratios of bacteria (Table 1). We gradually tried to reduce the concentration of Z1 and increase the concentration of its competitors. This forced Z1 to produce more antibiotics and there was stronger selection pressure on their development. At first, the ratio was 1:1 (100 µl Z1 and 100 µl *S. aureus* in liquid culture) and then the concentration of Z1 and *S. aureus* bacteria changed. Each passage was checked under a microscope with a magnification of 1000x using immersion oil. From the 16th passage, the temperature in the thermostat was increased to 14°C, then to 15°C for the 17th passage and then by 2°C each time if Z1 bacteria prevailed in the given passage. When a passage began to show resistant behavior *of S. aureus* against the cultured antibiotic, new passages were created from the original *S. aureus* from the first passage and the cultured Z1 bacteria were purified by cross-staining on an agar plate of a Petri dish, where individual colonies were isolated. A pure colony of Z1 bacteria was placed in a new passage with *S. aureus* from the first passage. Then the passage was monitored and new passages were formed. When the passages showed the functioning of the antibiotic and the dominance of Z1 bacteria over *S. aureus*, the temperature could be increased. Culture of the antibiotic with Z1 bacteria required a slower increase in temperature of around 28°C, and only by 1°C. At these temperatures, the antibiotic is probably more unstable. When testing the effectiveness of an antibiotic, the maximum temperature limit at which the antibiotic was cultured must be observed. Therefore, it is very appropriate to observe the specified temperature limit or lower the temperature by 1-2°C when testing antibiotics. The Production of the antibiotic was bred by gradually increasing the temperature up to 38°C.

**Table 1.** Cultivation and breeding of Z1 bacteria against *S. aureus*.

| Cultivation and breeding of Z1 bacteria and *Staphylococcus aureus* |  |  |
| --- | --- | --- |
| passage number | Concentration Z1 against *S. aureus* | temperature |
| 1.-8. | 1:1 | 12°C |
| 9. | 1:4 | 12°C |
| 10. | 1:5 | 12°C |
| 11. | 1:10 | 12°C |
| 12.-14. | 4:50 | 12°C |
| 15. | 5:100 | 12°C |
| 16. | 3:100 | 12-14°C |
| 17th-18th | 3:100 | 15°C |
| 19. | 1:10 | 15°C |
| 20. | 3:100 | 15°C |
| 21. | 1:100 | 15°C |
| 22nd-23rd | 1:100 | 18°C |
| 24. | 3:100 | 20°C |
| 25. | 1:10 | 20°C |
| 26. | 1:10 | 22°C |
| 27. | 1:10 | 22-24°C |
| 28. | 1:40 | 26-31°C (increasing by 1°C after 4-7 days) |
| 29. | 1:40 | 32°C |
| 30. | 1:40 | 33°C |
| 31. | 1:40 | 34°C |
| 32. | 1:40 | 35°C |
| 33. | 1:40 | 36°C |
| 34. | 1:40 | 37°C |
| 35. | 1:40 | 38°C |

The same procedure was used for *S. agalactiae* with a different start of cultivation at 22°C. *S. agalactiae* bacteria showed poor growth at lower temperatures (12-15°C), therefore cultivation was started only at 22°C. Bacteria from passage no. 26 against *S. aureus* were used for cultivation. The Z1 bacterium had not yet encountered the new *S. agalactiae* bacterium, but it mostly dominated in the cultures. The first passage was created in a 1:1 concentration ratio at 22°C. The cultures were cultivated on mixed LB agar medium with Tryptone-soy agar medium (medium 71). Subsequently, the temperature was increased by 1-2°C, only in the case of clear dominance of Z1 against *S. agalactiae* . The last passage was tested by diffusion disk test at 38°C. All subsequent passages from No. 2 to No. 5 were created in concentration ratios of 3:10.

The same procedure was used for the bacterium *A. baumannii* CCM 2355, when the cultured bacteria Z1 was used against *S. aureus* at 38°C. The cultures were grown on mixed LB agar medium with Tryptone-soy agar medium (medium 71). The first passage was created in a 1:1 bacterial concentration ratio at 38°C. The second to fifth passages were created in a 2:1 concentration ratio. An antibiotic produced by the cultured bacteria Z1 was isolated from passage no. 5 and tested by the diffusion disk method.

The same procedure was used for *E. coli*, where cultured Z1 bacteria, which had been competently challenged with *A. baumannii* at 38 °C, were mixed at 38 °C with *E. coli* on LB medium. The first passage was made at a 1:1 bacterial concentration ratio at 38 °C. The second to fifth passages were made at a 2:1 concentration ratio. The sixth and seventh were made at a 3:1 ratio for *E. coli*. The antibiotic produced by cultured Z1 bacteria was isolated from passage no. 7 and tested by the diffusion disk method.

Similarly for *P. aeruginosa*, when cultured Z1 bacteria, which was after a competitive fight with *E. coli* at 38 °C, was mixed at 38 °C with *P. aeruginosa*, on a mixed LB agar medium with tryptone-soy agar medium (medium 71) in a ratio of 1:1. The first passage was made in a ratio of 1:1 bacterial concentration at 38 °C. The second to fifth passages were made in a concentration ratio of 2:1. The sixth and seventh were in a ratio of 3:1 for *P. aeruginosa*. The antibiotic produced by cultured Z1 bacteria was isolated from passage no. 7 and tested by the diffusion disk method.

A similar procedure was used for the yeast *S. cerevisiae*. The first contact with the Z1 bacteria in the first passage was at 12°C and all other passages were in 1:1 concentration ratios. From passage number 4, the temperature was increased to 15°C and in passage number 6, the temperature was increased to 18°C. The passages with cell cultures were continuously monitored. Testing of bacterial breeding for the production of antibiotics against the yeast *S. cerevisiae* took half a year.

### Isolation of antibiotics

The antibiotic was isolated using an agar plate with antibiotic-producing bacteria, ethyl acetate and filter paper (for diffusion discs) (according to Horová 2022) (6). 65 µl of Z1 bacteria and 135 µl of *S. aureus* bacteria were inoculated onto 14 Petri dishes with a diameter of 90 mm containing LB agar. The bacteria were spread over the entire agar plate with a hockey stick and placed in a thermostat for 24 hours. At 12°C for the first isolation, at 22°C for the second isolation, at 32°C for the third isolation, and at 38°C for the fourth. After 24 to 48 hours, the grown bacterial cultures were scraped off and removed. The agar plates were then placed in a beaker with ethyl acetate (50ml). In ethyl acetate, the agar plates were crushed for better isolation of the antibiotic and the entire beaker was left for 24 hours on a shaker at room temperature. The entire beaker was covered with a transparent plastic membrane protecting the mixture from evaporation. After 24 hours, the remains of the agar plate were removed from the solution and filter papers intended to absorb the antibiotic were inserted into the solution. The entire mixture was allowed to evaporate at 55°C. The resulting filter paper discs were placed in the center of a new agar plate with bacteria of the species *S. aureus* (original uncultured culture) and placed in a thermostat for 24 hours and optionally at room temperature for 24 hours. As a control, the same filter paper discs were used, they were impregnated with ethyl acetate but without antibiotic and the ethyl acetate was allowed to evaporate. This is advantageous not only for creating the best possible control conditions, but also because the ethyl ester of acetic acid will destroy any bacterial contamination on the surface of the filter paper.

The same procedure was used for Z1 bacteria for competitive interaction against *S. agalactiae.* Z1 bacteria (30 µl) and *S. agalactiae* (100 µl) were plated on 30 Petri dishes with a diameter of 60 mm. Similarly, *A. baumannii* was plated on 30 Petri dishes with a diameter of 90 mm together with Z1 bacteria on mixed LB agar medium with Tryptone-Soy Agar (medium 71) at a rate of 100 µl *A. baumannii* and 30 µl Z1. Similarly, antibiotics against *E. coli* and *P. aeruginosa* were obtained. The mixed cultures were placed in a thermostat at 38°C for 24 hours.

Sequencing of the 16S rRNA region using the Illumina method using ViennaLab techniques was performed at Bioptická laboratoř s.r.o. (Mikulášské náměstí 628/4, 326 00 Czech Republic).

## Results

### Species identification

When attempting to identify the bacterium designated as Z1, sequencing of the 16S rRNA Illumina MiSeq (V3–V4) region found a match with *Pantoea* sp. 57917 82.31% at readcount (% of classified reads). The highest match is with *Pantoea agglomerans* according to the SILVA database. However, the sequenced region has a high BLAST match with other species, e.g. *Lelliottia* sp. strain SWJTUDBP9, *Enterobacter* sp., *Klebsiella michiganensis* strain RHB20-C02, *Klebsiella pneumoniae* strain ARLG-3185 chromosome, *Leclercia adecarboxylata* strain M17, *Salmonella enterica* subsp. *enterica* serovar Dublin strain CVM N53043 chromosome, *Pectobacterium wasabiae* NAPoBL202 and so on. Therefore, the studied bacterium cannot be unambiguously assigned to the species *P. agglomerans*. The presence of several other sequence variants with 1–3 nucleotide differences and their BLAST matches with other species of the genus *Pantoea* (e.g. *P. anthophila*, *Pantoea* sp. DAFNE) also suggest that the isolate may represent a previously undescribed species within this genus. This variability may be a consequence of the natural diversity of multiple copies of the 16S rRNA gene or amplification artifacts. Given these findings and the unique phenotypic properties of the isolate, such as the production of new antibiotics, it is quite likely that this may be a new species with potential for biotechnological applications. Further taxonomic analyses, including multilocus sequence analysis (MLSA) and whole-genome sequencing, will be necessary to confirm this hypothesis. It is also necessary to note that the currently used strain has probably undergone a number of mutations due to long breeding and may already differ in a number of genes from the original wild strain.

### Staphylococcus aureus

When Z1 bacteria first came into contact with *S. aureus* bacteria in the first passage, there were no signs of antibiotic production. Even after several days, the ratios of the bacteria were balanced (Fig. 2). Gradually, during subsequent passages at a concentration of 1:1 at 12°C, Z1 bacteria began to prevail over *S. aureus.* This is evidenced by passage number 8, where Z1 bacteria prevailed over *S. aureus* by an estimated 95% after just 4 days (Fig. 3). The passages were continuously monitored and their development was monitored even after a longer time interval, in liquid cultures sometimes even after 2 months. From the 9th passage, the concentrations changed from 1:1 to 1:4, when the amount of *S. aureus* was increased and, conversely, the amount of Z1 bacteria decreased. However, after 1 day of competitive interaction, the number of Z1 and *S. aureus* equalized to a ratio of 1:1. This is a sign of antimicrobial production against *S. aureus*. A tailored antibiotic against *S. aureus* can be obtained as early as passage 8. It is ideal to let the bacteria interact for more than 3 days, as the Z1 bacteria have more time to produce antibiotics and suppress competing *S. aureus* bacteria.

**Fig. 2.**
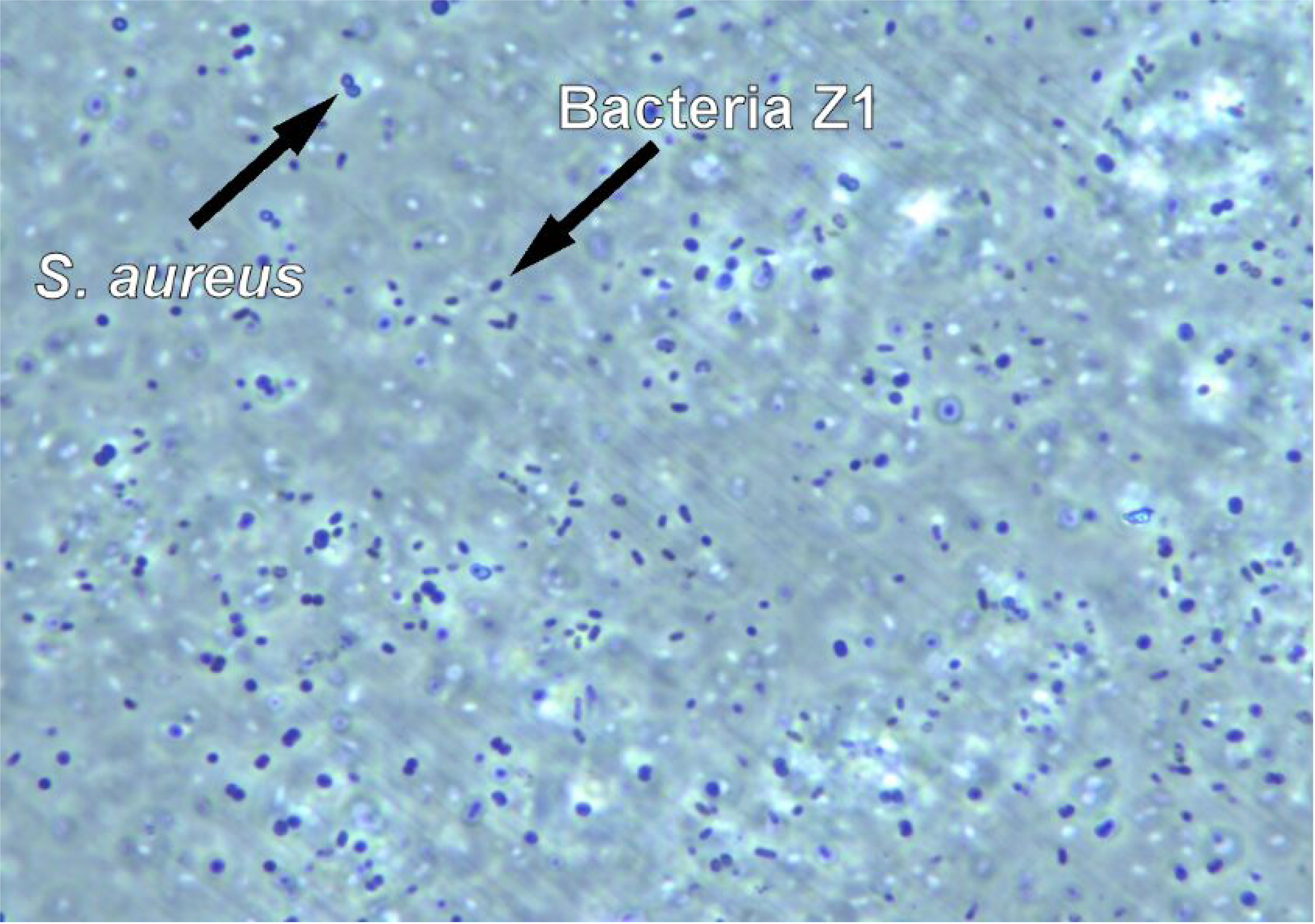
Passage 1 with *S. aureus* and Z1 bacteria after 9 days in liquid LB medium, concentration 1:1 at 12°C (magnification 1000x).

**Fig. 3.**
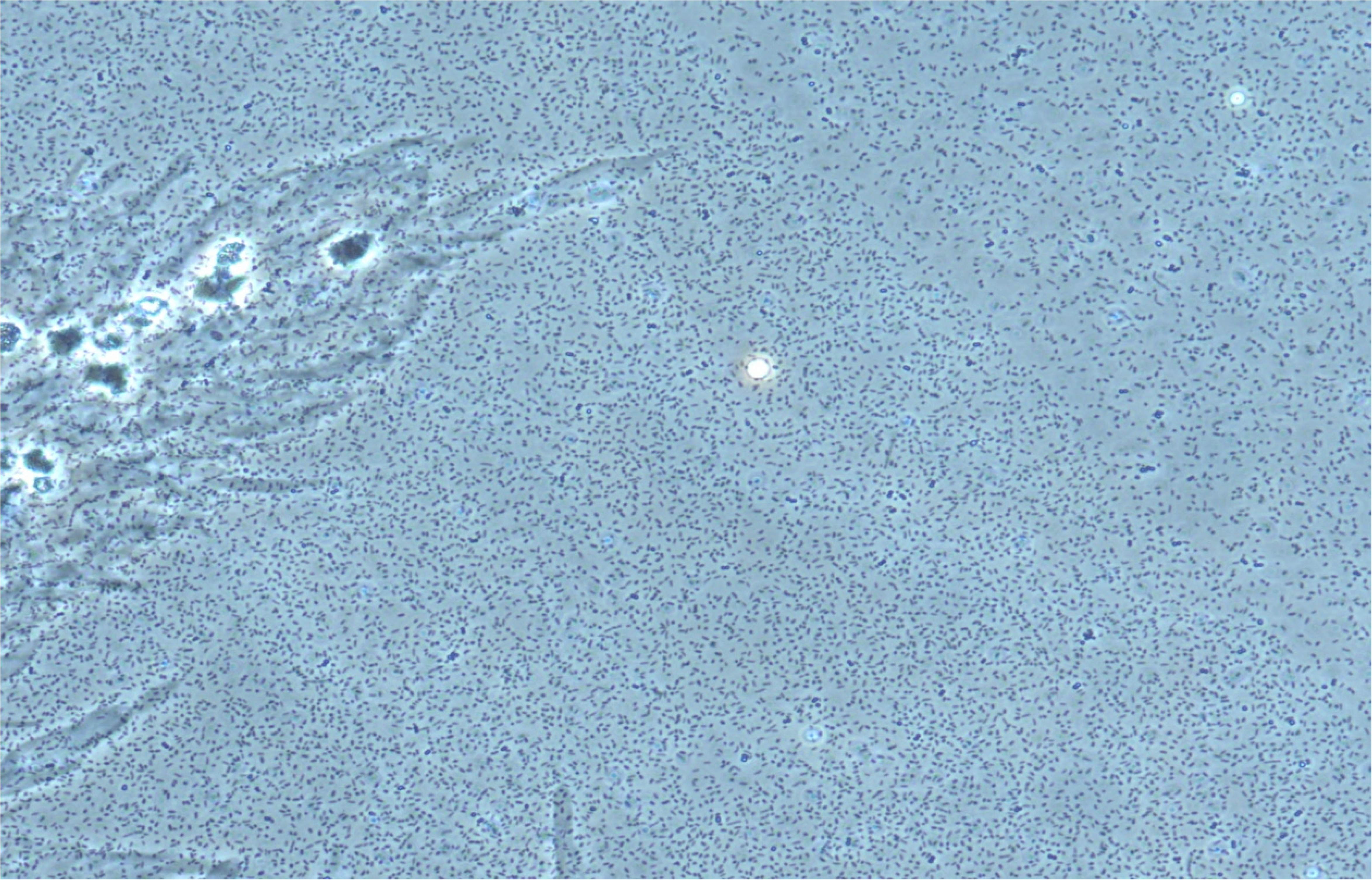
Passage No. 8 after 4 days of competitive interaction between Z1 and *S. aureus* bacteria in a 1:1 ratio of liquid LB medium at 12°C. Numerical superiority of Z1 bacteria over *S. aureus* after 4 days (magnification 1000x).

To provide clear evidence of antibiotic production by Z1 bacteria against other bacteria, a diffusion disk test was performed on agar plates of Petri dishes. In the first phase of cultivation, the produced antibiotic was isolated from the 12th passage, which was cultivated at 12°C. The experiment was performed on two Petri dishes, one at 12°C and the other at room temperature. *S. aureus* bacteria from the first passage, when this bacterium had not yet encountered a new antibiotic, were seeded on the Petri dishes. Subsequently, antibiotic soaked in filter papers was placed in the centers of the Petri dishes. The result was the formation of an inhibition zone around the filter paper, where the isolated antibiotic against *S. aureus* bacteria was present. However, this inhibition zone was only formed on the Petri dish that was stored in a thermostat at 12°C (Fig. 4). The second Petri dish stored at room temperature did not contain any sign of an inhibition zone (Fig. 5).

**Fig. 4.**
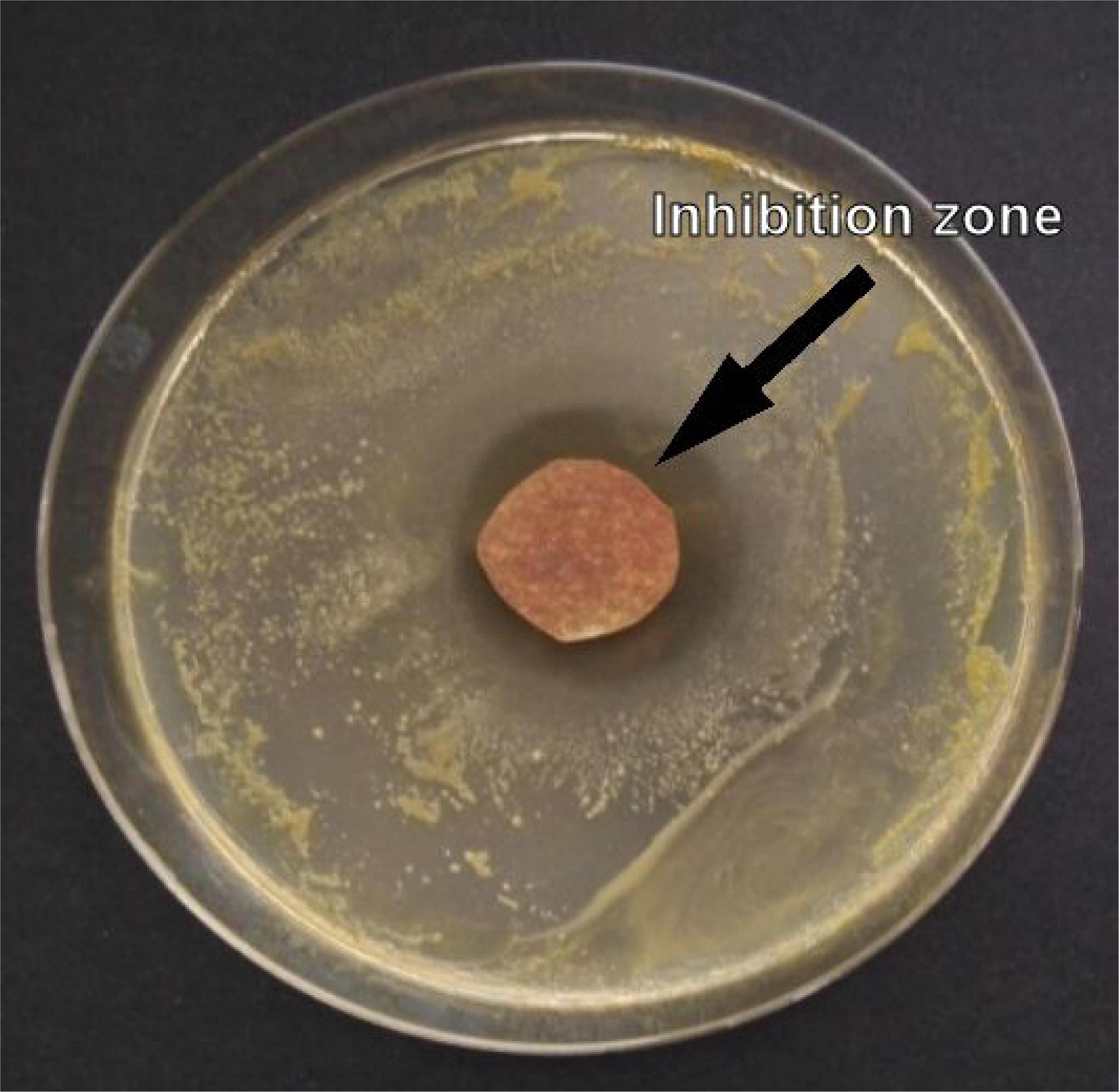
Disk diffusion test of the isolated antibiotic of the bacterium Z1 from agar on filter paper against *S. aureus* at 12°C with inhibition zone.

**Fig. 5.**
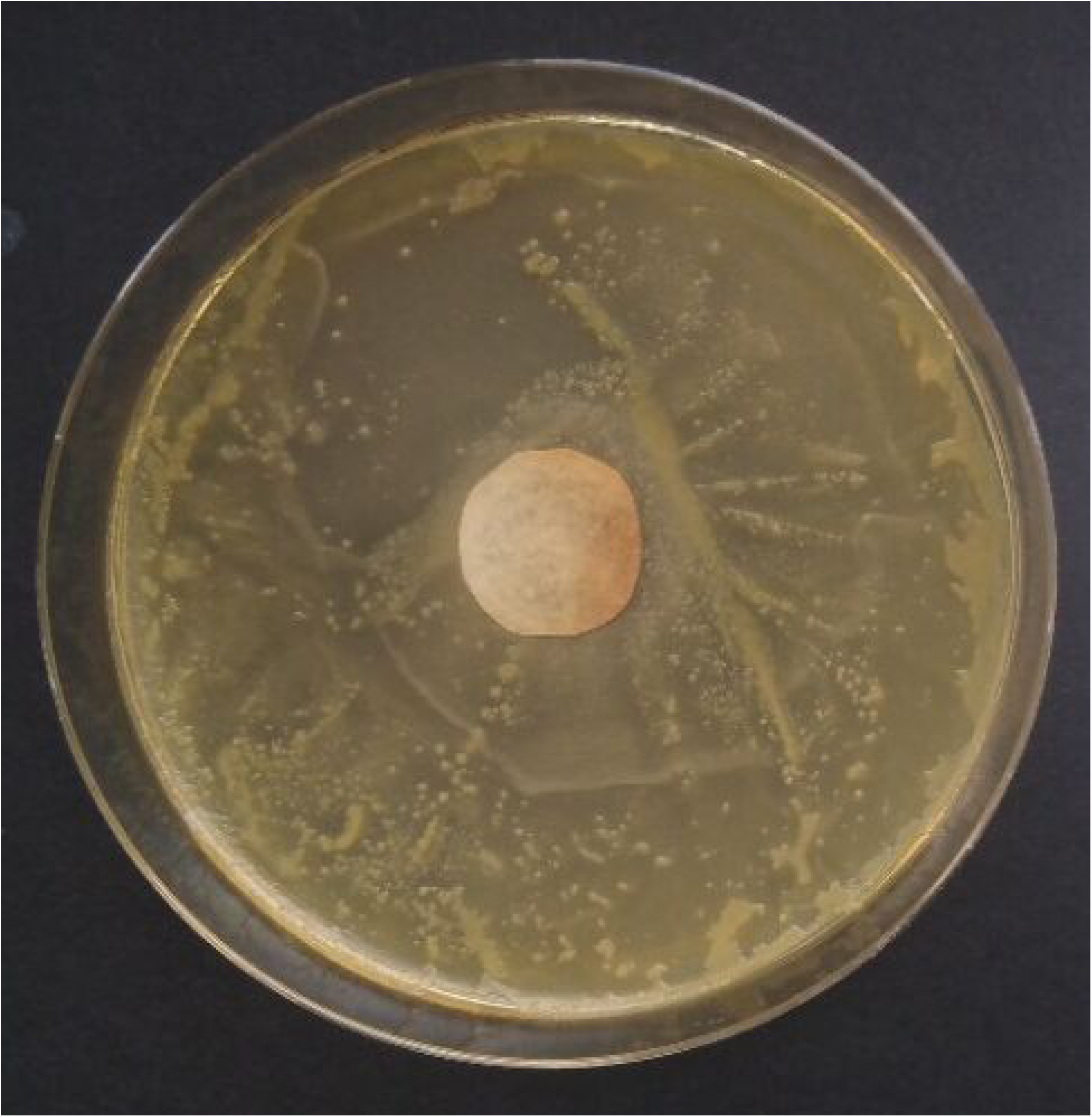
Disk diffusion test of the isolated antibiotic of the bacterium Z1 from agar on filter paper against *S. aureus* at room temperature without an inhibition zone.

Since the inhibition zone did not form in the Petri dish at room temperature, the antibiotic with Z1 bacteria was cultured to function and grow at higher temperatures than only 12°C. However, before increasing the temperature, it was necessary to adapt the Z1 bacteria so that it could suppress many *S. aureus* in a small amount of Z1. For example, in the 15th passage, the concentration of Z1 bacteria against *S. aureus* was given in a ratio of 5:100. Passage number 15 was cultured at 12°C in liquid LB medium on a shaker, when 50 µl of Z1 bacteria and 1000 µl of *S. aureus* bacteria were given. After 24 hours, the result of the superiority of Z1 bacteria over *S. aureus* was very significant (Fig. 6). Similarly, in the 8th passage, the Z1 bacteria again gained numerical superiority over *S. aureus*.

**Fig. 6.**
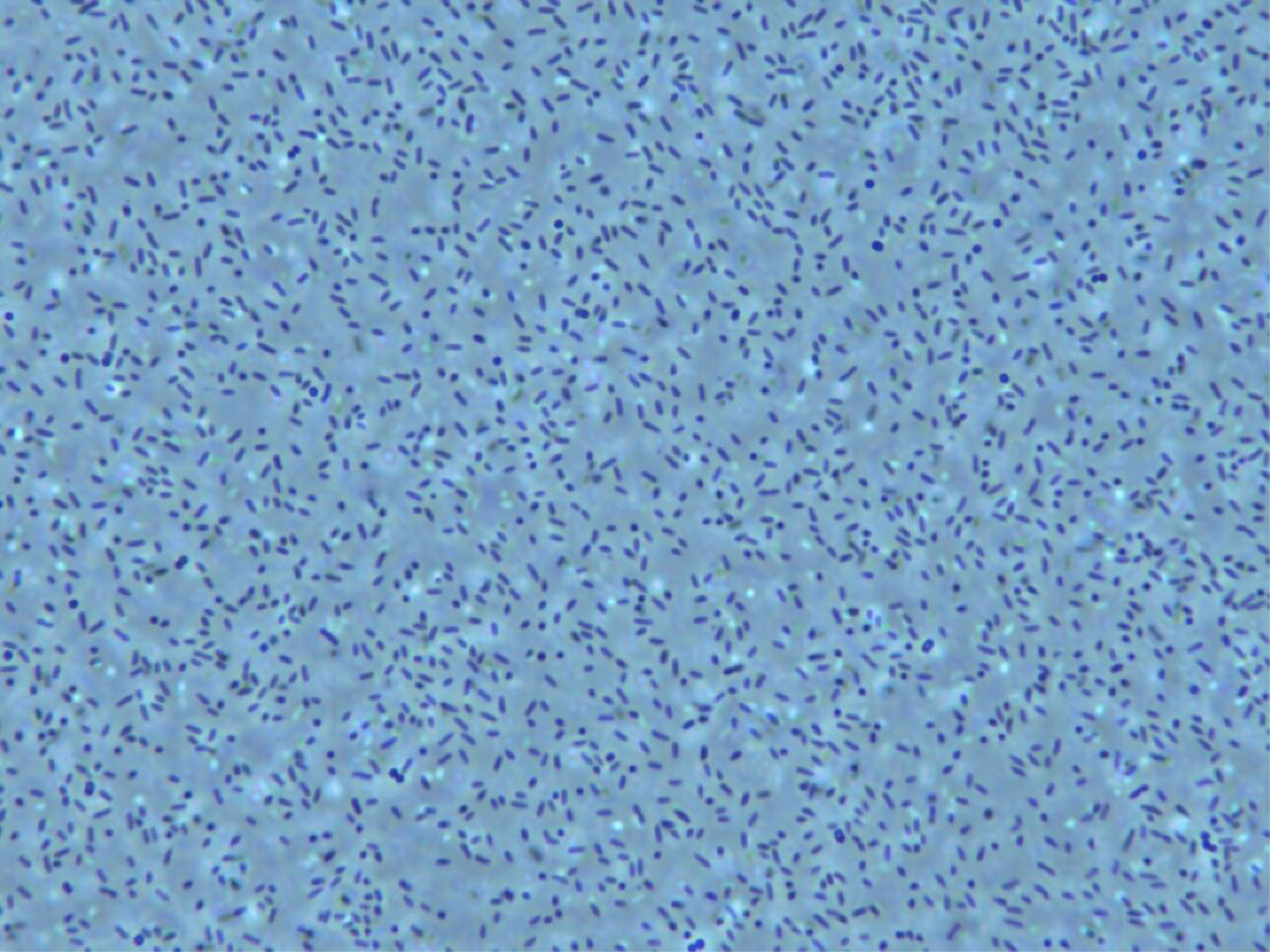
Passage no. 15 of competitive interaction between Z1 and *S. aureus* bacteria in a ratio of 5:100 of liquid LB medium at 12°C. Numerical superiority of Z1 bacteria over *S. aureus* after 24 hours (magnification 1000x).

Even after several days, there was a clear predominance of Z1 bacteria over *S. aureus bacteria* in the 15th passage. Z1 bacteria dominated the sample with an estimate of approximately 98%. A similar result was recorded in the 16th passage.

Because the results of the cultivation of the 15th passage and the 16th passage were very satisfactory, the temperature was increased from 12°C to 14°C by 2°C for the 16th passage. However, the concentration of bacteria was maintained and the 17th passage was also created in a concentration ratio of 3:100 (30 µl Z1 and 1000 µl *S. aureus*). The temperature was gradually increased by two degrees for each passage. Each passage was monitored continuously for several days. Only cultures where the Z1 bacteria had significantly established themselves were selected for further cultivation (Fig. 7).

**Fig. 7.**
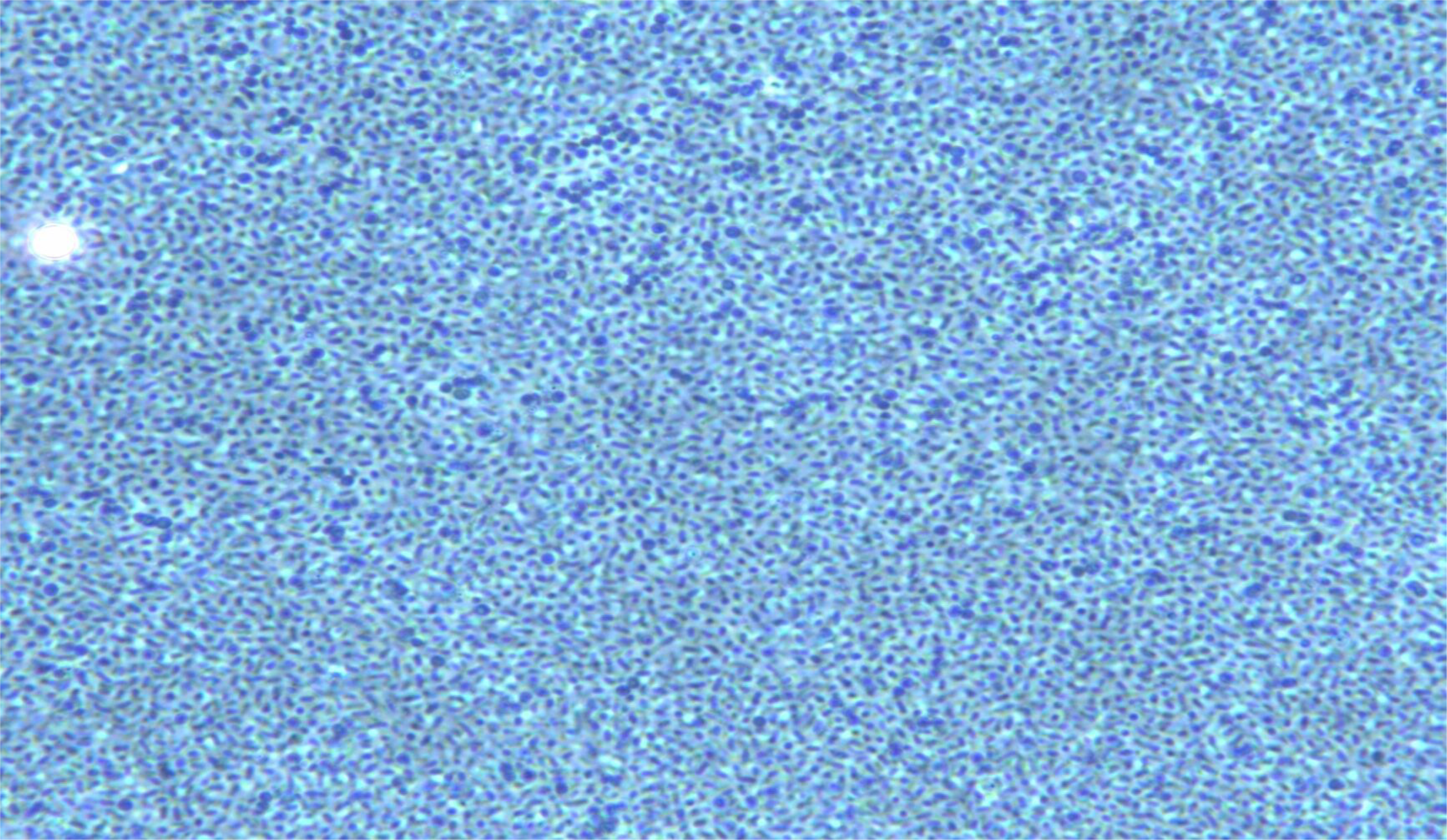
Passage no. 26 of competitive interaction between Z1 bacteria and *S. aureus* in a ratio of 1:10 on agar medium at 22°C. Numerical superiority of Z1 bacteria over *S. aureus* after 4 days (magnification 1000x).

The results of the inhibition zone formation were also checked after 3 days. After 3 days, the inhibition zones had decreased. This may be due to the selection pressure on *S. aureus* bacteria, so some could mutate to migrate to other parts of the Petri dishes where there are more nutrients. Another reason may be the instability of the isolated antibiotic, which may break down after several days. However, after checking the diffusion disk test after 2 weeks, the inhibition zone was still clear on the Petri dishes and the antibiotic still had some effect even after 2 weeks.

Fig. 8 shows the final experiment with antibiotics against *S. aureus.* This is a 34 passage with Z1 bacteria grown at 38 °C. Antibiotics were obtained as in previous cases using ethyl acetate from 25 plates, where Z1 and *S. aureus* bacteria competed against each other at a ratio of 3:10 (30 µl Z1 and 100 µl *S. aureus*).

**Fig. 8.**
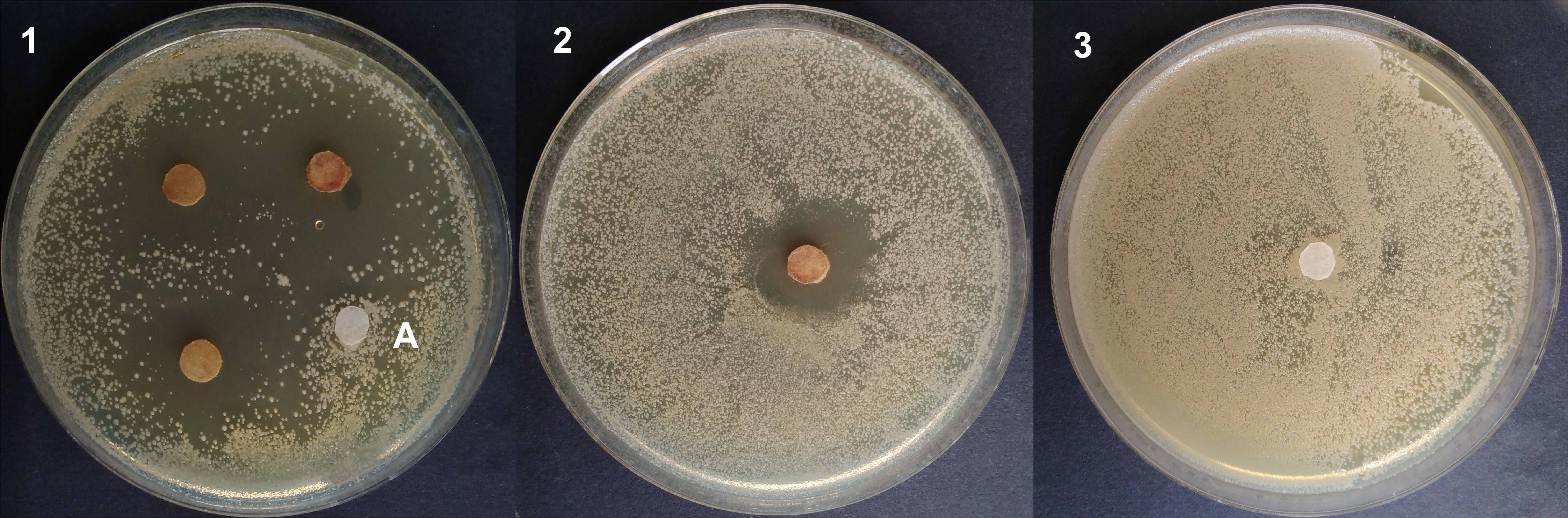
Disk diffusion test of the isolated antibiotic of the bacterium Z1 from agar on filter paper against *S. aureus* at 38 °C. 1 - disks with antibiotic, disk marked A is the control 2 - separate disk with antibiotic, the width of the inhibition zone was about 2.5 cm, 3 - control only, disk without antibiotic.

### Streptococcus agalactiae

The breeding of the antibiotic against *S. agalactiae* was carried out similarly to that for *S. aureus*. For the new experiments, the Z1 against *S. aureus* cultured from higher temperatures was used, specifically the Z1 bacteria cultured at 38°C. These bacteria adapted very quickly to the first new contact with the *S. agalactiae* bacteria and already in the 4th passage very promising results were obtained (Fig. 9). The advantage was mainly their ability to grow at higher temperatures than the temperatures around 12-20°C. The breeding process was greatly accelerated in this way. Passage number 4 was cultured at a temperature of 38°C in a ratio of 3:10. It showed the best breeding results and suppressed most of the *S. agalactiae* bacteria. The *S. agalactiae* bacteria without contact with the Z1 bacteria grow very well at 38°C and show no signs of suppression (Fig. 10).

**Fig. 9.**
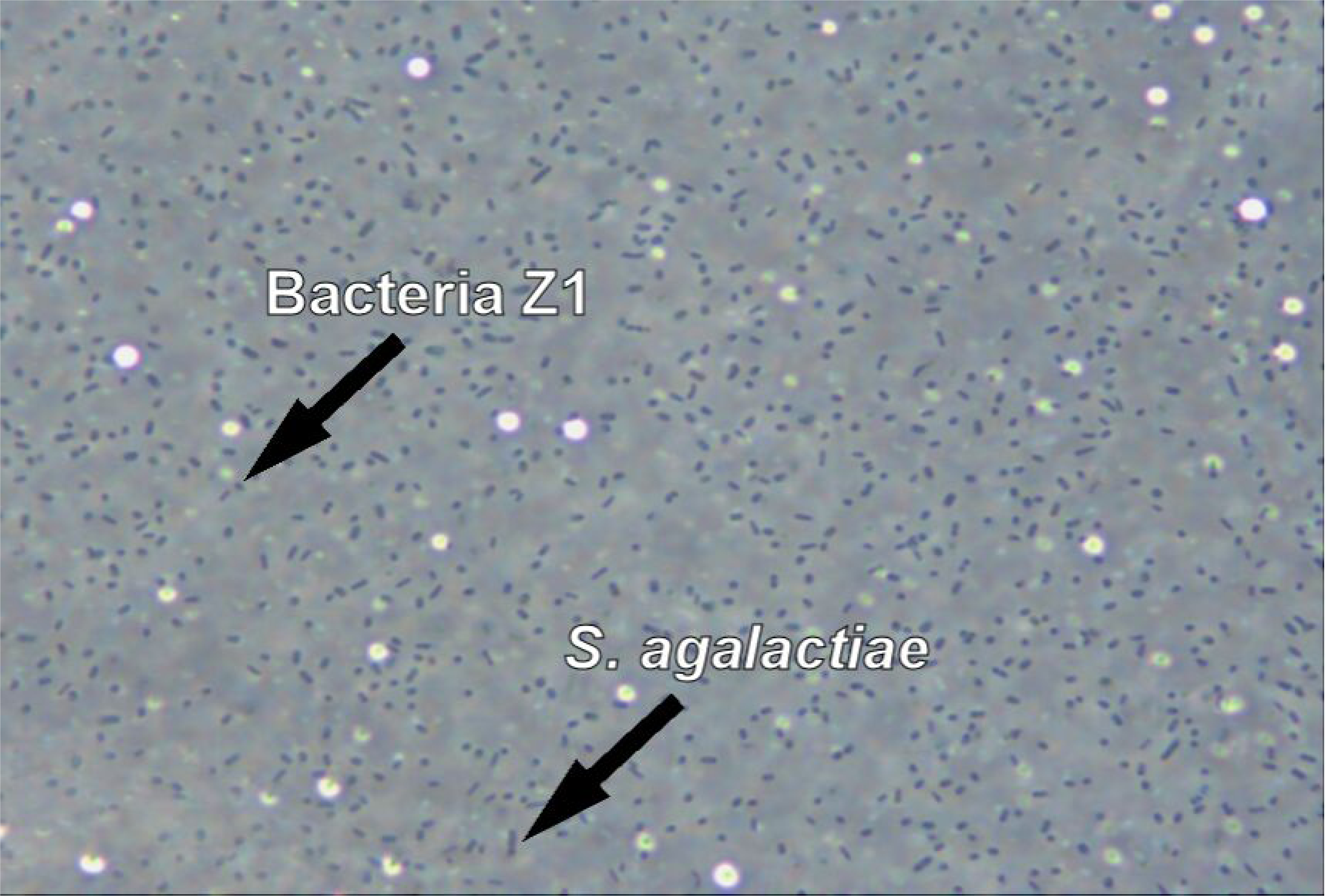
Passage no. 4 of competitive interaction between bacteria Z1 and *S. agalactiae* in a ratio of 3:10 on agar medium at 30°C. Numerical superiority of bacteria Z1 over *S. agalactiae* after 4 days (magnification 1000x).

**Fig. 10.**
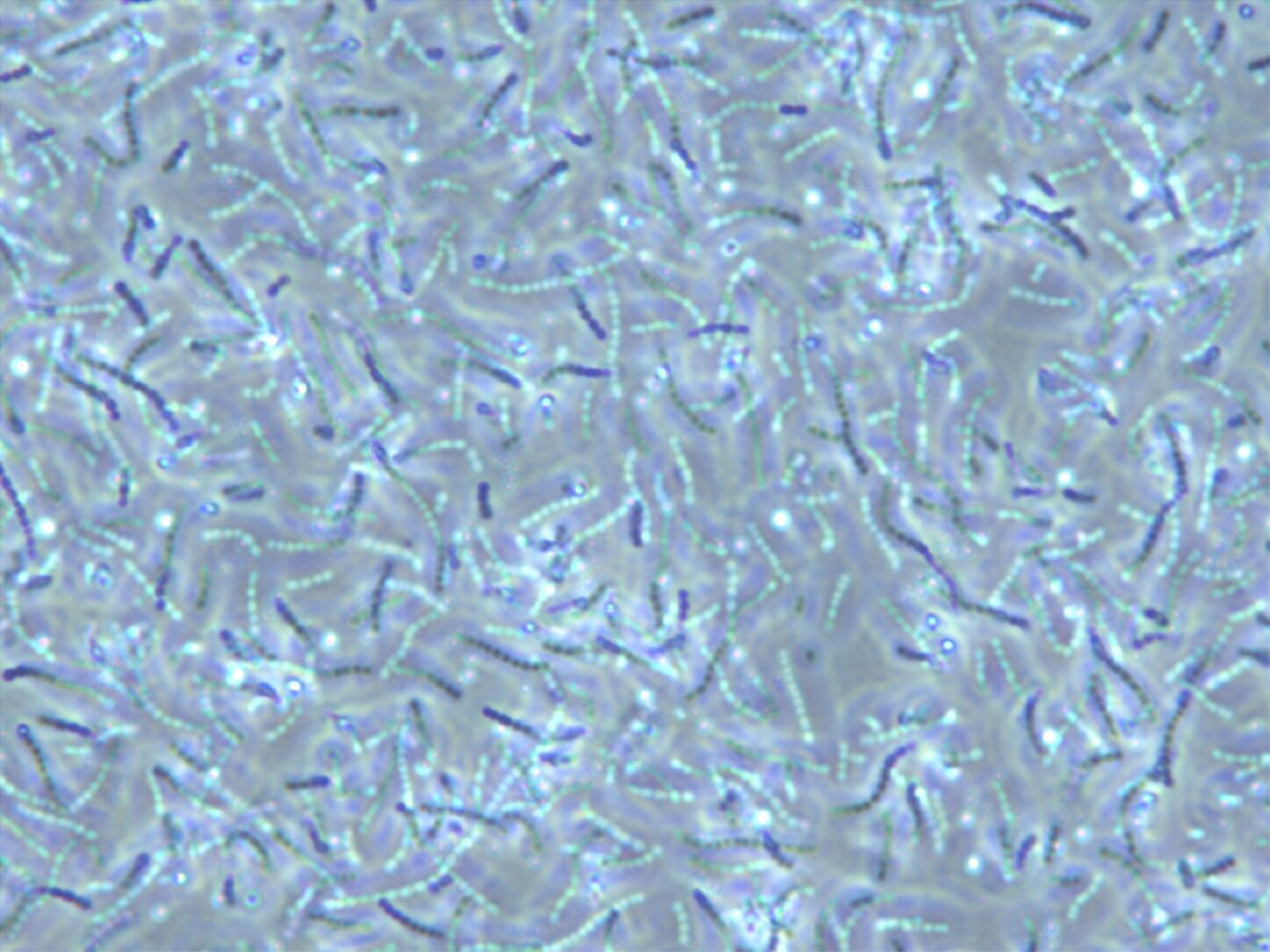
Pure culture of *S. agalactiae* in agar medium at 38°C without contact with Z1 bacteria after 3 days (magnification 1000x).

A 5th passage was created, which was used to create a relatively strong concentration of the antibiotic. The antibiotic was obtained from forty 90 mm diameter dishes at 38°C, where Z1 and *S. agalactiae* competed at a ratio of 3:10 (Fig. 11). Even after 10 days, the antibiotic was still effective against *S. agalactiae* in the Petri dish, but the inhibition zone gradually decreased.

**Fig. 11.**
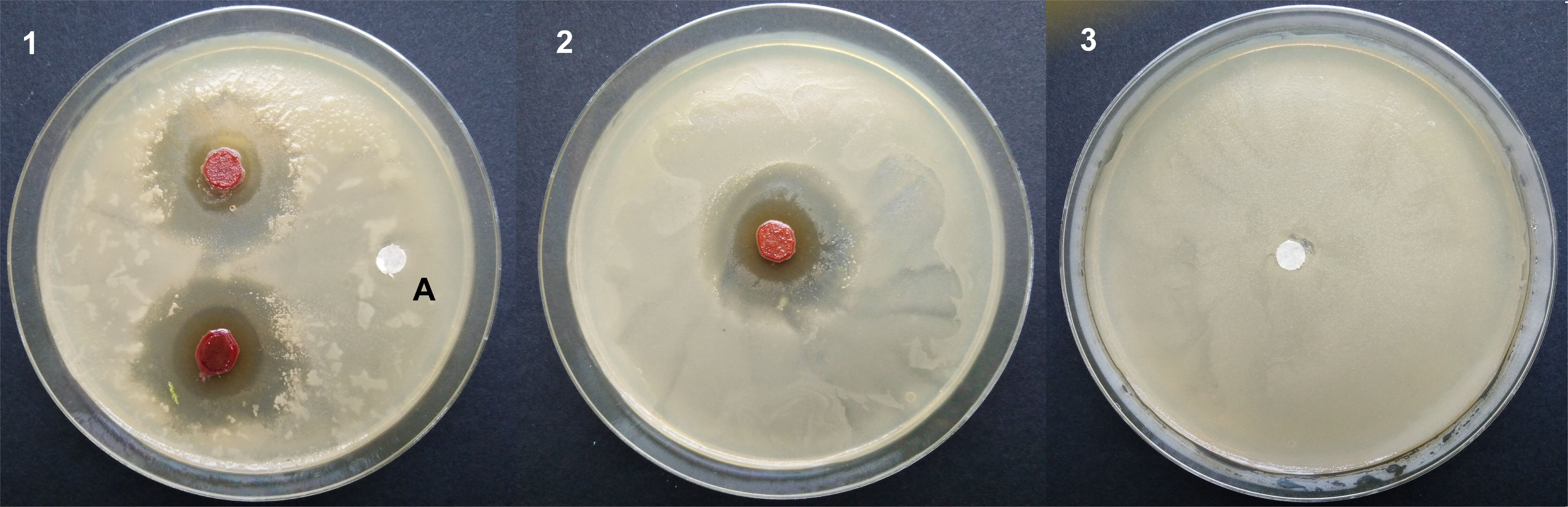
Disk diffusion test of the isolated antibiotic of the bacterium Z1 from agar on filter paper against *S. agalactiae* at 38°C. 1 - disks with antibiotic, disk marked A is the control 2 - separate disk with antibiotic, the width of the inhibition zone was about 2 cm, 3 - control only, disk without antibiotic.

### Acinetobacter baumannii

The breeding of the antibiotic against *A. baumannii* proceeded similarly to that for *S. aureus* and *S. agalactiae*. Breeding began at 38°C on agar in a Petri dish at a concentration ratio of 1:1, but after three to five days Z1 began to prevail. In subsequent passages Z1 began to dominate more quickly and efficiently (Fig. 12). The antibiotic was obtained from forty 90 mm diameter dishes at 38°C, where Z1 and *A. baumannii* competed at a ratio of 3:10 (Fig. 13).

**Fig. 12.**
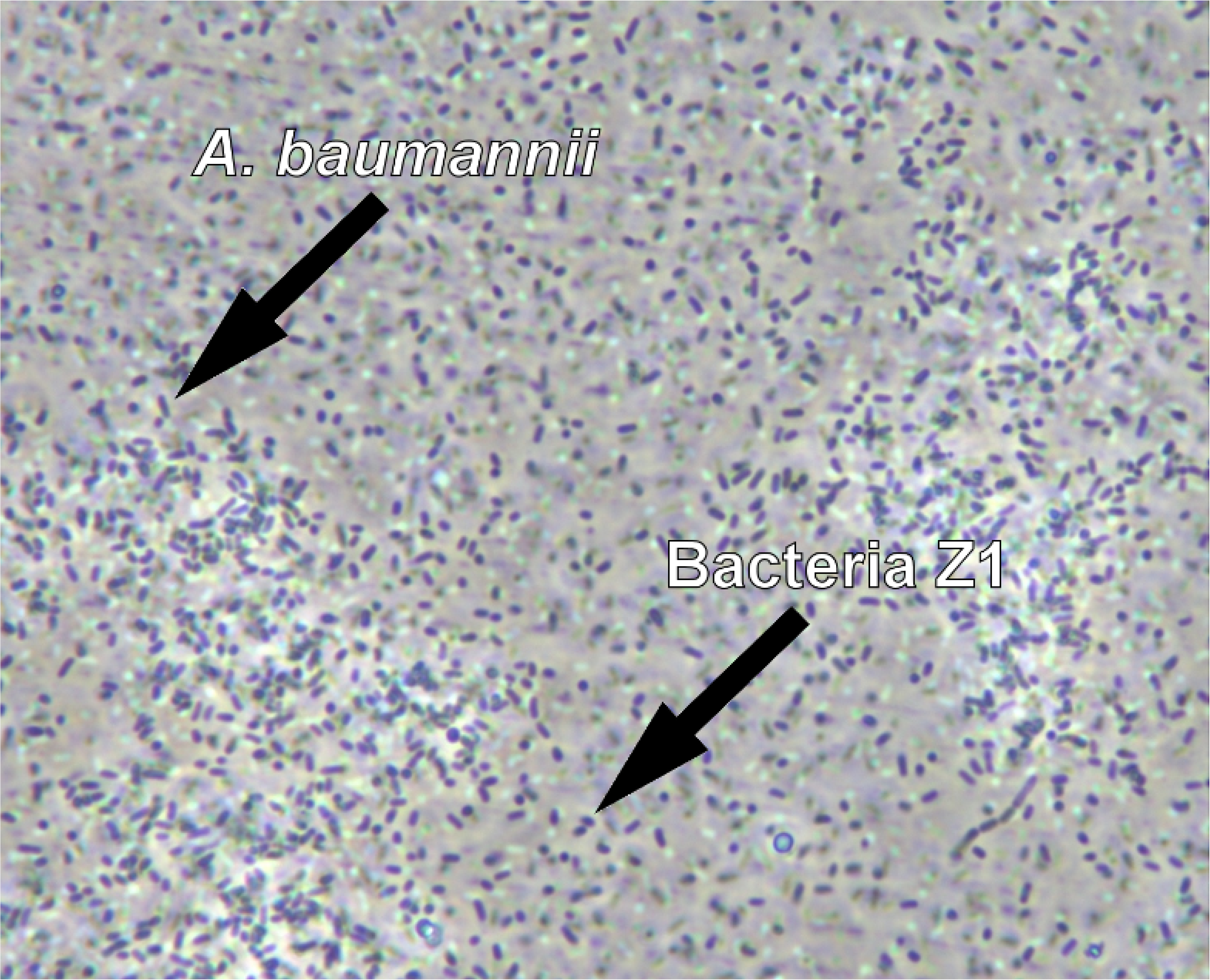
Passage no. 4 of competitive interaction between Z1 bacteria and *A. baumannii* in a ratio of 2:10 at 38°C (magnification 1000x) for antibiotic isolation.

**Fig. 13.**
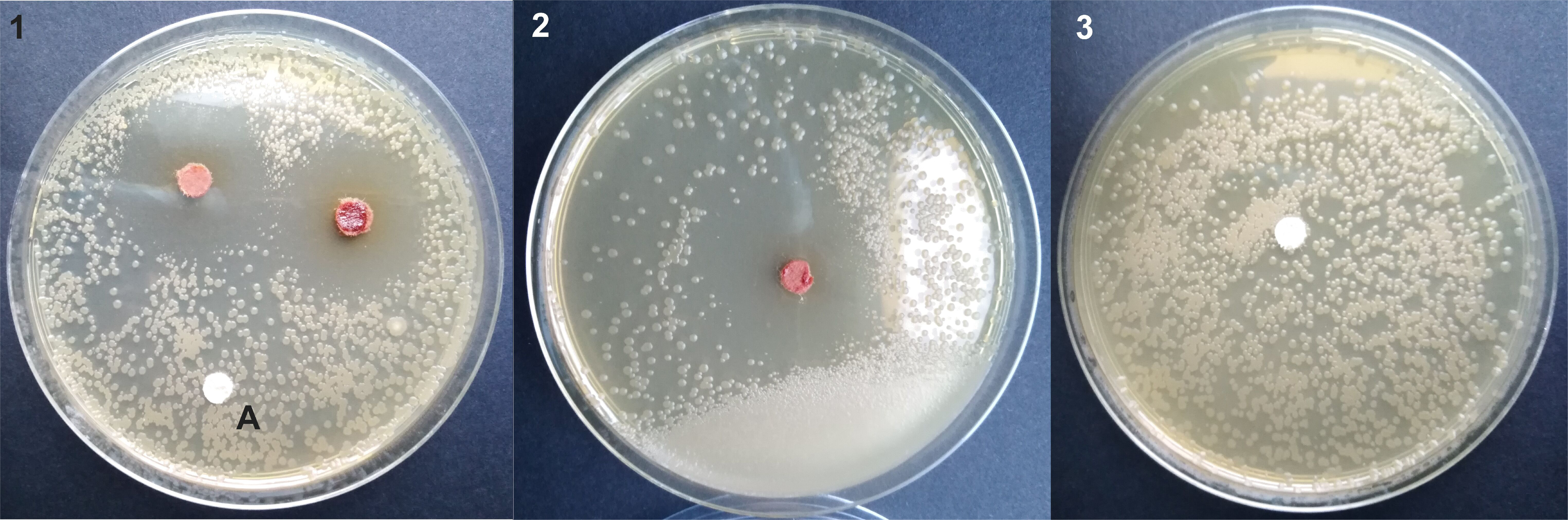
Disk diffusion test of the isolated antibiotic of the bacterium Z1 from agar on filter paper against *A. baumannii* at 38°C. 1 - disks with antibiotic, disk marked A is the control 2 - separate disk with antibiotic, the width of the inhibition zone was about 2 cm, 3 - control only, disk without antibiotic.

### Escherichia coli

The breeding of the antibiotic against E. coli proceeded similarly to that against *A. baumannii.* The competitive fight against *E. col*i, however, was more fierce than with other bacterial opponents. It took more passages before the Z1 bacteria began to win, but even then some *E. coli* always survived. Probably, during the breeding, the *E. coli* was also bred to become a resistant strain. Only in the dish where the seven-day competitive fight took place and the agar in the dish began to dry out, only Z1 survived, and they then won more easily in contact with the non-resistant E. coli strain (Fig. 14). An antibiotic was then obtained from these Z1 (Fig 15). The antibiotic was obtained from forty 90 mm diameter dishes at 38°C, where Z1 and *E. coli* competed at a ratio of 3:10 (Fig. 15).

**Fig. 14.**
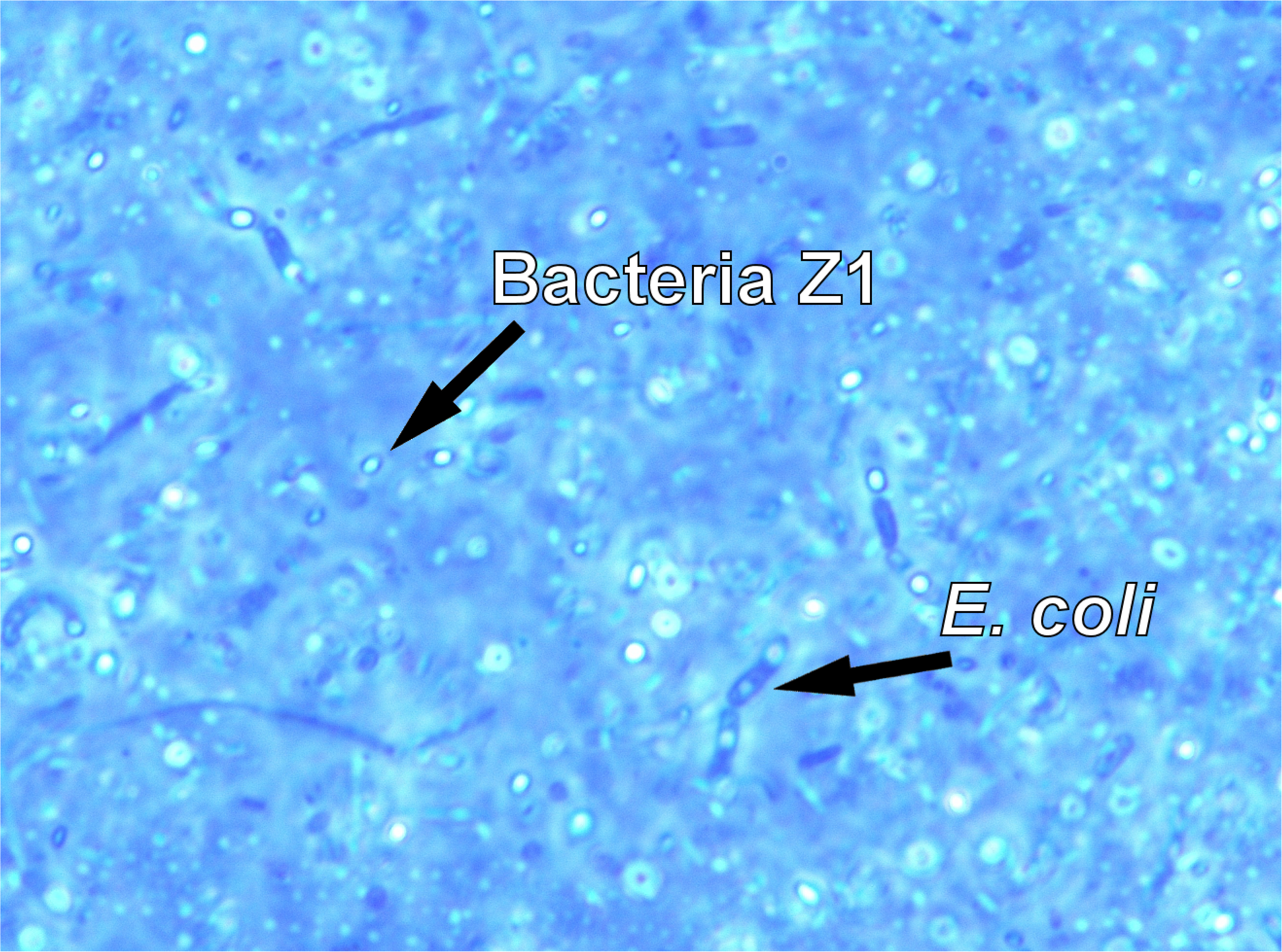
Passage no. 7, Z1 bacteria and *E. coli* after 4 days, in a ratio of 3:10, at 38 °C (magnification 1000x).

**Fig. 15.**
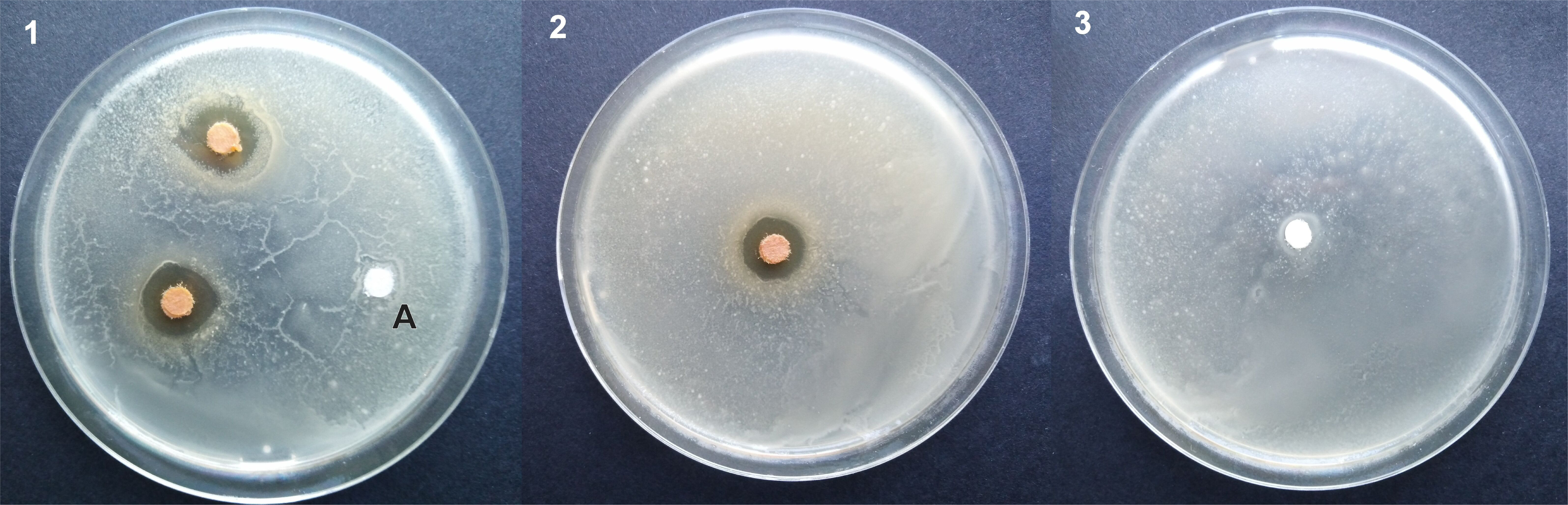
Disk diffusion test of the isolated antibiotic of the bacterium Z1 from agar on filter paper against *A. baumannii* at 38°C. 1 - disks with antibiotic, disk marked A is the control 2 - separate disk with antibiotic, the width of the inhibition zone was about 1.4 cm, 3 - control only, disk without antibiotic.

### Pseudomonas aeruginosa

The cultivation of the antibiotic against *P. aeruginosa* was carried out similarly to that against *E. coli*. The concentration of Z1 against *P. aeruginosa* was gradually reduced and seven passages were carried out (Fig. 16). In this type of cultivation, the competitive struggle always took a relatively long time, only after 5-6 days did the Z1 bacteria gain a clear advantage. The antibiotic was then obtained from the Z1 bacteria from the seventh passage in forty 90 mm diameter dishes at 38 °C, where Z1 and *P. aeruginosa* competed in a ratio of 3:10 (Fig. 17).

**Fig. 16.**
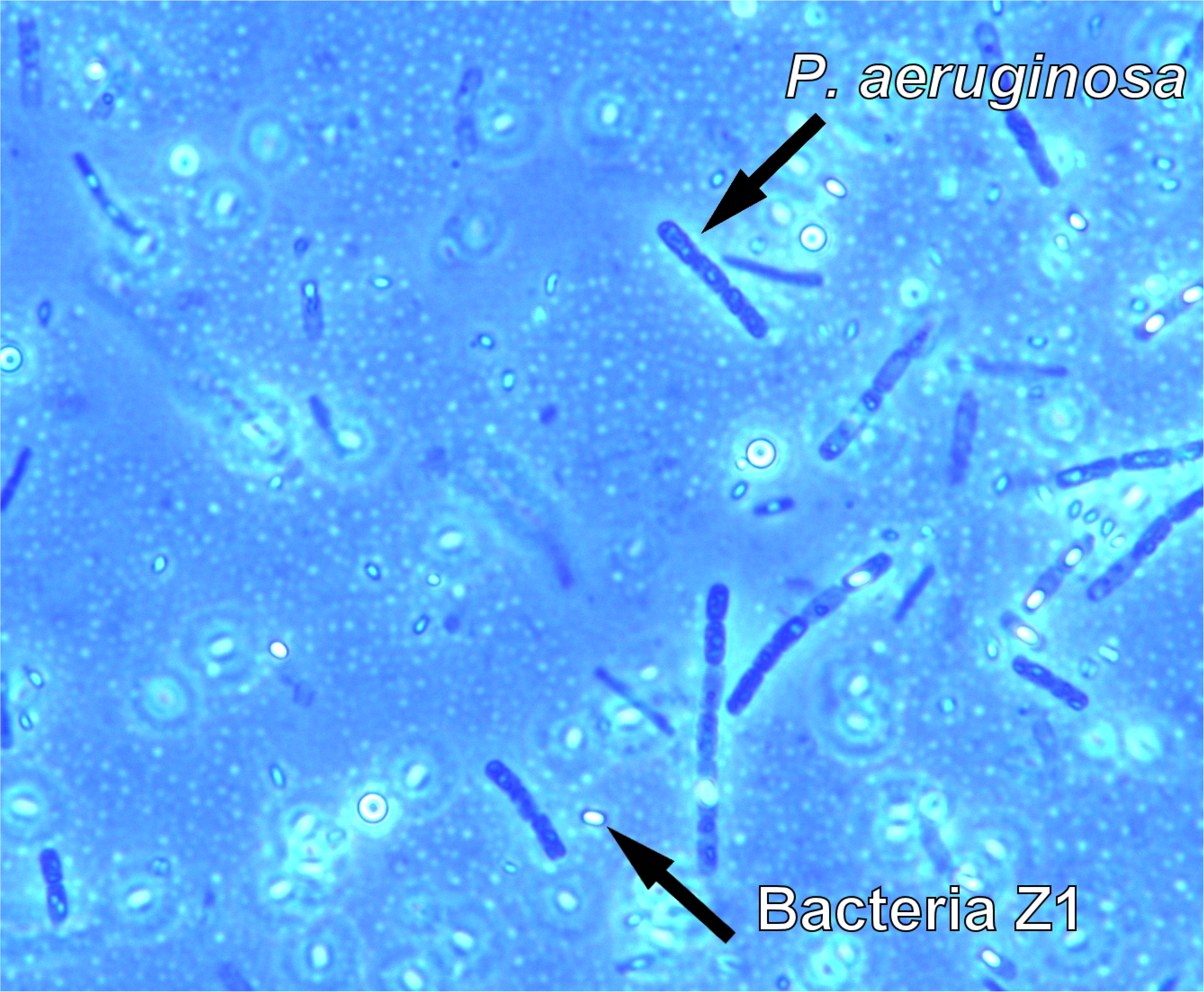
Passage No. 1, Z1 bacteria and *P. aeruginosa* after 4 days, in a ratio of 1:1, at 38 °C (magnification 1000x).

**Fig. 17.**
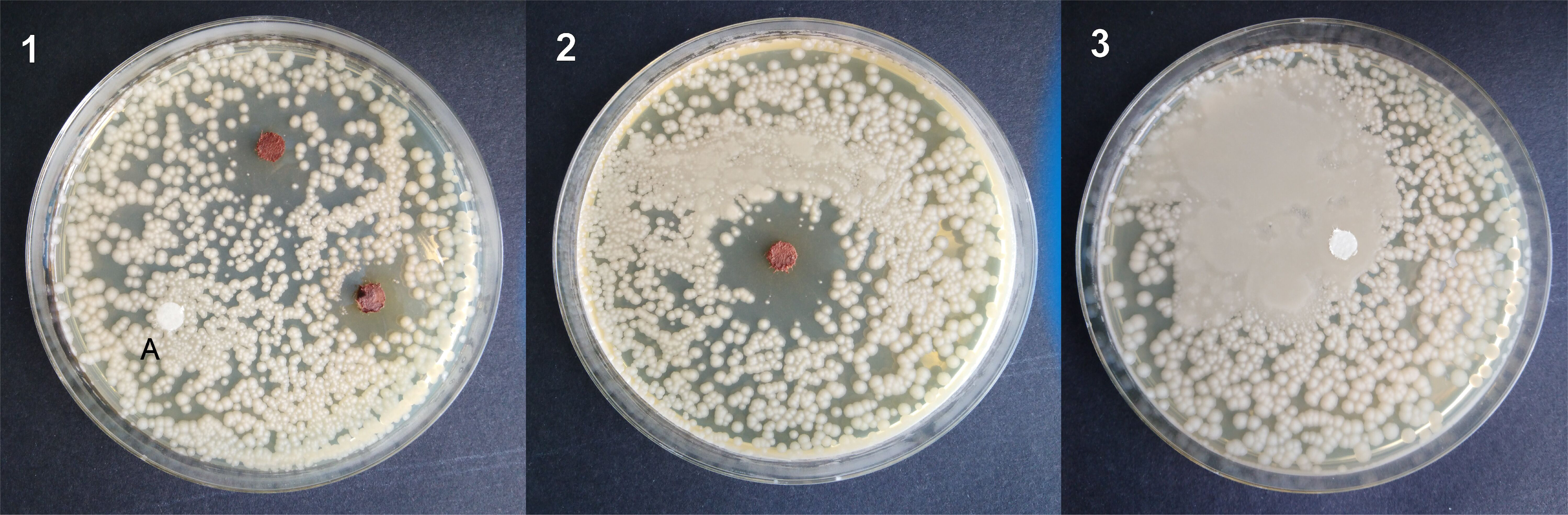
Disk diffusion test of the isolated antibiotic of the bacterium Z1 from agar on filter paper against *P. aeruginosa* at 38 °C. 1 - disks with antibiotic, disk marked A is the control, the width of the inhibition zone was about 1.5 cm, 2 - a separate disk with antibiotic, the width of the inhibition zone was about 2 cm, 3 - control only, disk without antibiotic.

### Saccharomyces cerevisiae

An attempt to produce antibiotics was also carried out on *S. cerevisiae*. The cultivation of antibiotics began at 12°C in a concentration ratio of 1:1 and the temperature was gradually increased as with *S. agalactiae* and *S. aureus*. After the second contact of Z1 bacteria with *S. cerevisiae yeast*, there was no production of antibiotics and suppression of the yeast (Fig. 18). Even in passage number 4, which was placed in a thermostat at 15°C, there were no signs of antibiotic production against *S. cerevisiae* yeast (Fig. 19). The cultivation of antibiotics and the overall production of antibiotics by Z1 bacteria apparently does not work against *S. cerevisiae* yeast, since Z1 bacteria did not numerically prevail over the yeast.

**Fig. 18.**
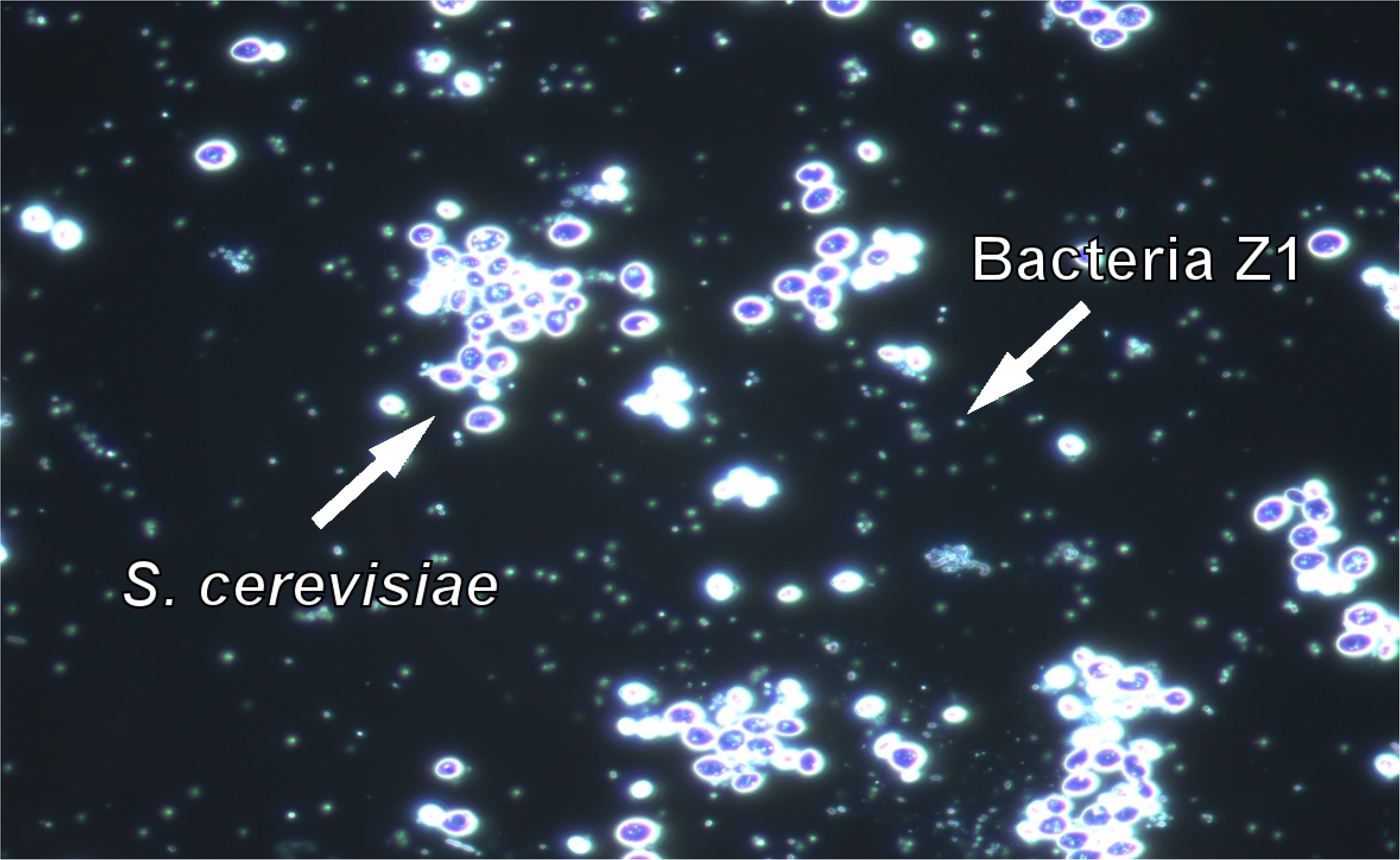
Passage number 2 with Z1 bacteria and S. cerevisiae yeast after 7 days in liquid LB medium, concentration 1:1 at 12°C (magnification 1000x). No signs of antibiotic production against *S. cerevisiae* yeast.

**Fig. 19.**
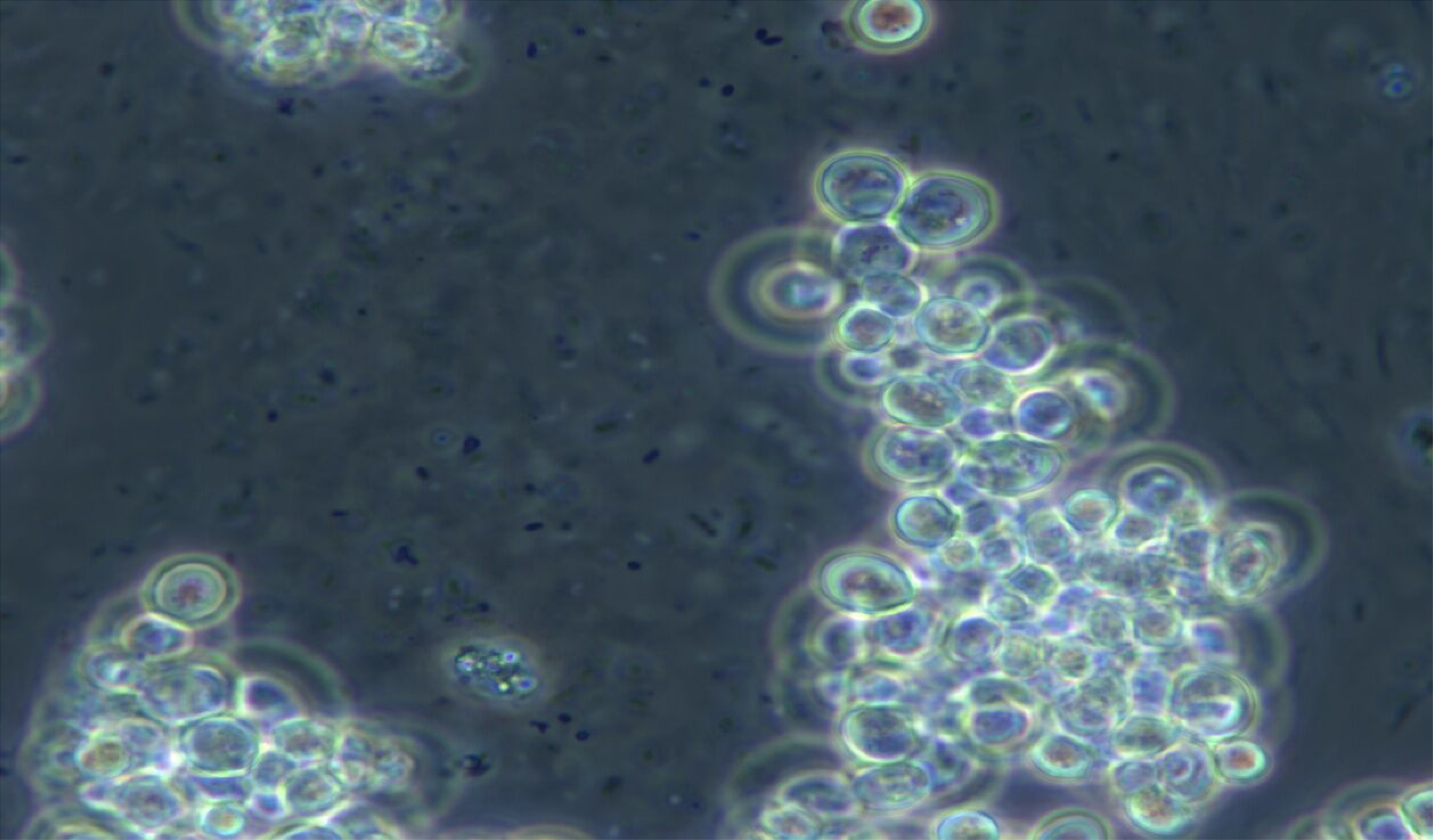
Passage number 4 with Z1 bacteria and S. cerevisiae yeast after 21 days in liquid LB medium, concentration 1:1 at 15°C (magnification 1000x). No signs of antibiotic production against *S. cerevisiae* yeast.

## Discussion

Our data support a robust and cost-effective method to induce antibacterial activity in a non-genetically modified bacterial strain through ecological pressure. Importantly, this methodology does not require advanced screening technologies or molecular tools, making it suitable for low-resource research environments.

At this stage of research, obtaining new types of antibiotics through biological means seems to be a relatively easy matter. However, there are still many unclear facts. We will focus on the positives first. The Z1 bacterium is capable of producing antibiotics against other types of bacteria. We do not need genetic manipulations, we just need to naturally breed bacteria, rely on random mutations and take into account natural evolutionary trends. Obtaining strains that produce new antibiotics is relatively very fast and cheap. Another advantage is that the new antibiotics are apparently relatively narrowly focused on one strain and do not affect other types of bacteria. The molecules of these antibiotics are probably not very stable, so they could be easily broken down in the body.

The negatives include that we have to rely on chance, we do not know what molecules are being created and that the resulting antibiotics are unlikely to be very stable compounds, which could be a problem during storage.

At first glance, it is clear that the most effective antibiotic is against *S. aureus*, which is clear evidence of a long breeding process. There were 35 passages against *S. aureus*, while against other bacteria it was only 5-7 passages. This is a clear message that if we need a stronger antibiotic, the breeding process needs to be made appropriately longer.

Nowadays, gene manipulations are very popular, which are very useful but financially expensive. Conventional breeding for example for disease resistance is tedious, time consuming and mostly dependent on environment as compared to molecular breeding, particularly marker assisted selection, which is easier, highly efficient and precise (9). However, the evolution and breeding of bacteria to produce antibiotics is less expensive and more naturally imitates evolution in nature. The advantage of evolutionary breeding of bacteria is speed.

Other authors have also explored methods for improving antibiotic-producing strains, including experimental evolution and genome shuffling to enhance compound diversity and potency (10–12). In contrast, our method represents a form of targeted breeding through direct interbacterial competition, where the selective pressure is not artificially imposed but emerges from ecological interactions.

Bacteria have the great advantage of rapid reproduction and the evolution of their generations, which also adapt the antibiotics they produce. Of course, bacteria will develop resistance to these new antibiotics. It was remarkable to watch how *E. coli* resisted even during breeding and created resistant forms while Z1 produced increasingly effective antibiotics, and so even though Z1 won, some *E. coli* were already resistant when they were finally in the minority against Z1. But Z1 bacteria and similar ones will probably be able to create new antibiotics against resistant strains in the future.

Based on the results, it seems very likely that it will be possible to produce different antibiotics against a specific type of bacterial disease by breeding bacteria. Such antibiotics should not burden the body by suppressing other types of bacteria that naturally live in the human body. The antibiotic would be narrowly specialized only for a specific type of bacteria. However, it could also work on closely related bacteria, but that could be a new topic for further study.

While the chemical identity of the inhibitory compounds remains unknown, their selective activity and lack of effect on *Saccharomyces cerevisiae* suggest specificity and limited eukaryotic toxicity. This raises the potential for further medical, veterinary, or agricultural applications.

Future work will include the characterization of antibiotic molecules via chromatography and mass spectrometry, cytotoxicity assays on mammalian cells, and full genome sequencing of Z1. Our method may also be extended to other bacterial genera, enabling broader discovery of ecologically evolved antibacterial phenotypes.

Therefore, further research should focus on testing the bred antibiotics on eukaryotic cells, especially mammals, and finally, it is necessary to examine the effect on the human body. However, it is not yet clear whether these antibiotics will not harm mammalian cells, which is probably the most important. It would also be very appropriate to try to breed antibiotics, for example, using the proven bacterium Z1 against other types of bacteria and potential pathogens. This seems to be a relatively long road. But there is hope that a new antibiotic produced in this way could be quickly produced and tested against some plant pathogens. For example, the bacterium *Xylella fastidiosa* is a major pest that attacks a large number of plant species and causes significant damage. Furthermore, this pathogen is divided into a number of subspecies (13), so it would be quite advantageous to produce specific antibiotics against each subspecies.

There are also other unresolved questions, it is necessary to map the genome of the Z1 bacterium and determine this bacterium to the exact species. For practical use, it is necessary to characterize the antibiotic molecules. It seems quite likely that Z1 produces a different type of antibiotic against each competitor. The Z1 bacterium probably belongs to the genus *Pantoea,* therefore it can be assumed that the antibiotic molecules will be similar to the pantocin A molecule, pantocin B, PNP-3 a PNP-4 and herbicolin O. The pantocin A molecule is acid, base and thermal lable (8)(14). It is also likely that the molecules also differ depending on the temperature at which a particular strain functions. It was very surprising to find that antibiotic molecules produced at 12 °C do not function at room temperature, and therefore it was necessary to breed the strain of this rather cold-loving bacterium so that it could function well at human body temperature. It would be very disadvantageous if the bacteria produced antibiotics, for example, at 30 °C and they would then not function at 37 °C. Therefore, it would also be appropriate to characterize the produced antibiotic molecules at different temperatures and compare them with each other. Based on the experiments performed, it is likely that Z1 creates different types of antibiotic molecules depending not only on the competing species, but also on the temperature.

## Conclusions

This study presents a practical and reproducible methodology to evolve antibiotic-producing bacteria through interspecies competition and selective passage. The approach yielded a single strain of *Pantoea* with clear inhibitory effects against multiple clinically relevant pathogens. The method is immediately applicable for laboratories seeking new bioactive strains without access to high-throughput chemical screening. Continued development and refinement may help accelerate the discovery of novel antibacterial agents.

## ACKNOWLEDGMENTS

I would like to thank the Centre for Biology, Geosciences and the Environment, Faculty of Education, University of West Bohemia, from whose institutional funds the project was financed. We acknowledge the BC CAS core facility LEM supported by the Czech-BioImaging large RI project (LM2023050 and OP VVV CZ.02.1.01/0.0/0.0/18_046/0016045 funded by MEYS CR) for their support with obtaining scientific data presented in this paper.

